# Influence of multi-axial dynamic constraint on cell alignment and contractility in engineered tissues

**DOI:** 10.1101/2020.08.12.248039

**Authors:** Noel H. Reynolds, Eoin McEvoy, Juan Alberto Panadero Pérez, Ryan J. Coleman, Patrick McGarry

## Abstract

In this study an experimental rig is developed to investigate the influence of tissue constraint and cyclic loading on cell alignment and active cell force generation in uniaxial and biaxial engineered tissues constructs. Addition of contractile cells to collagen hydrogels dramatically increases the measured forces in uniaxial and biaxial constructs under dynamic loading. This increase in measured force is due to active cell contractility, as is evident from the decreased force after treatment with cytochalasin-D. Prior to dynamic loading, cells are highly aligned in uniaxially constrained tissues but are uniformly distributed in biaxially constrained tissues, demonstrating the importance of tissue constraints on cell alignment. Dynamic uniaxial stretching resulted in a slight increase in cell alignment in the centre of the tissue, whereas dynamic biaxial stretching had no significant effect on cell alignment. Our active modelling framework accurately predicts our experimental trends and suggests that a slightly higher (3%) total SF formation occurs at the centre of a biaxial tissue compared to the uniaxial tissue. However, high alignment of SFs and lateral compaction in the case of the uniaxially constrained tissue results in a significantly higher (75%) actively generated cell contractile stress, compared to the biaxially constrained tissue. These findings have significant implications for engineering of contractile tissue constructs.

## 1 Introduction

A growing interest in the biomechanical behaviour of cells seeded in 3D culture has emerged in recent years. Mechanical priming strategies have been developed in an ongoing drive to engineer tissues with increased functional viability (Baker et al., 2008; Berry et al., 2003; Billiar et al., 2005; Cummings et al., 2004; Isenberg and Tranquillo, 2003; Mauck et al., 2003, 2000; Seliktar et al., 2000). Previous studies demonstrate that the application of 3D substrate stretch can up-regulate gene expression, promote differentiation, cause cell orientation redistribution, increase proliferation, and increase extracellular matrix synthesis (Berry et al., 2003; Campbell et al., 2007; Foolen et al., 2014; Gabbay et al., 2006). Given that cells actively respond to loading in the 3D microenvironment, it is important to develop a fundamental mechanistic understanding of the effects of mechanical conditioning on 3D synthetic tissue constructs. Therefore, the development of a robust system that can apply different uniaxial and biaxial deformation regimes to engineered hydrogel constructs while measuring the active and passive force, as presented in the current study, can provide new insights into the link between tissue constraint and active force generation.

Stretching of 2D substrates containing semi-confluent cell monolayers reveals that stress fibres (SFs) exhibit stretch avoidance (Barron et al., 2007; Kaunas et al., 2005; Neidlinger-Wilke et al., 2001; Wang et al., 2001). Stretch avoidance has also been reported in 3D substrates (Foolen et al., 2014). However, it has been more commonly reported that SF orientations remain unchanged during cyclic deformation (Foolen et al., 2012; Gauvin et al., 2011a; Nieponice et al., 2007a; Wakatsuki and Elson, 2002; Wille et al., 2006; Zhao et al., 2013). Cell contractile forces have previously been measured in tissues subjected to uniaxial stretching (Wagenseil et al., 2004; Wakatsuki et al., 2001, 2000; Wille et al., 2006; Zhao et al., 2014, 2013). Biaxial stretching has been employed to characterize cellular and scaffold fibres deformations (Gilbert et al., 2006, Courtney et al., 2006, Stella et al., 2008). However, due to the significant technical challenge of tissue force measurement during dynamic biaxial stretching, the influence of constraint on dynamic force generation has not been reported to date. A study by Thavandiran et al., (2013) provides a quantitative comparison of cell alignment and stress fibre formation on biaxially and uniaxially statically constrained collagen gels. However, force measurement is not performed, so the influence of constraint and dynamic loading on cell contractility is not uncovered. The current study provides a key advance investigating the influence of tissue constraint and dynamic loading on cell contractility and cell alignment.

In this investigation a bespoke experimental system is developed for measurement of cell and tissues forces during biaxial and uniaxial dynamic stretching. In addition to measuring the evolution of cell and tissue force, cell alignment throughout biaxial and uniaxial tissues is also characterised. In order to interpret experimental measurements for uniaxially and biaxially loaded tissues, a novel thermodynamically motivated model for cell contractility and remodelling is combined with an anisotropic hyperelastic model of the collagen gel to simulate the uniaxial and biaxial experiments. Simulations accurately predict the alignment and contractility of cells in biaxial and uniaxial constrained tissues and reveal that actively generated cell stress is significantly higher in the case of uniaxial loading.

## 2 Materials and Methods

### 2.1 Sample Preparation

Human cardiomyocytes isolated from a 24 year old male donor were obtained from Promocell (C-12810 (lot number 4051303.1), Heidelerg, Germany). Cells were grown as per Promocell protocols. Cell passages between P8 and P12, with a split ratio of 1:2, were used for all experiments (cells were observed to become senescent at P14). One week after plating in T25 flasks cell monolayers were observed to reach 100% confluency. Cells were observed to be highly elongated and aligned with neighbouring cells. However, no evidence of myotube formation or spontaneous beating was observed. Therefore, the cells are considered to exhibit an immature cardiomyocyte morphology and biomechanical characteristics. Before creating the collagen-cell solution, Teflon moulds, stainless steel hangers, and raisers are autoclaved and prepared in ethanol cleaned UV-sterilised petri dishes. The base of the petri dishes are coated with a siliconising reagent to inhibit cell adhesion (Sigmacote, SL2, Sigma–Aldrich Ireland Ltd., Arklow). Collagen-cell solution is made using 10X phosphate buffered saline (PBS), 1 M Sodium Hydroxide (NaOH), 1 mg/mL collagen solution, and cells suspended in standard growth media supplemented with an additional 10% foetal calf serum (FCS). The final solution contains a 0.2mg/mL collagen concentration with 1×10^6^ cells/mL and is at physiological pH and ionic strength (Figure 1-A). Appropriate volumes are made so that the initial height, *H*0, of the collagen-cell solution in the mould is 3 mm (Figure 1-B). Once the solution is added to the mould, the sample is carefully placed in an incubator (37°C, 5% CO2, and 90% humidity) for 1 hour to form a gel. A sufficient amount of HCM growth media is added to the petri dish to ensure the gel is completely submerged. The entire mould, which contains the gelled collagen-cell solution, is raised slightly from the surface of the petri dish using stainless steel raisers. This prevents adhesion of the gel to the petri dish and ensures that media is free to flow beneath the mould. Therefore, the cell laden gel is fully engulfed with growth media during incubation. After 2 days of incubation, cell mediated tension results in gel contraction (Figure 1-C). These contractile gels are referred to as “tissues”, or specifically as “uniaxial tissues” or “biaxial tissues”. Further samples are created using no cells. They are fabricated using the same moulds and quantities of PBS, collagen, and media as that used to create tissue samples. These samples are gelled by incubating at 37° for 1 hour. These cell free constructs are hereon referred to as “uniaxial hydrogels” or “biaxial hydrogels”.

**Figure 1:**
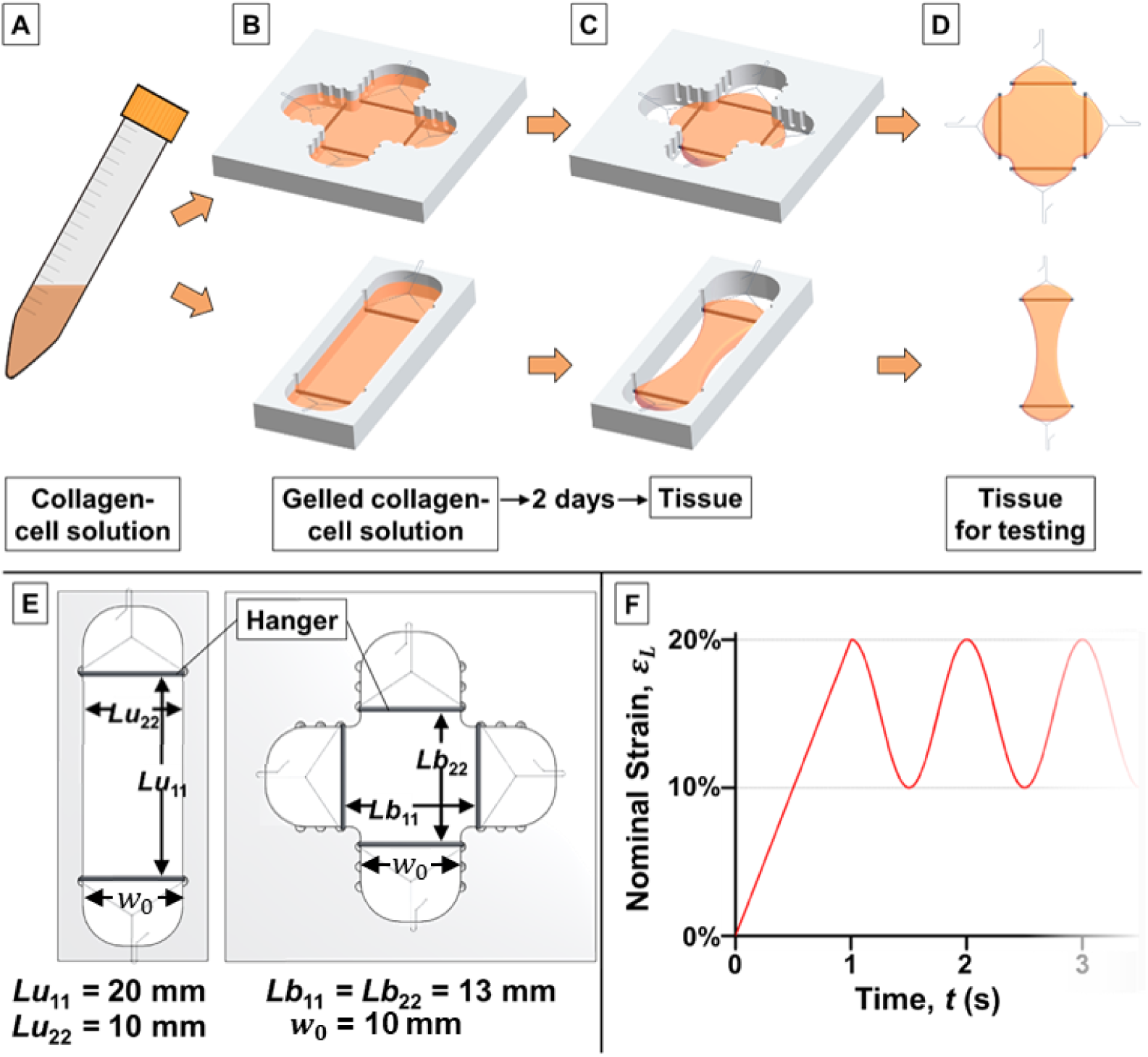
Schematic representation of sample preparation. (A) Collagen-cell solution is prepared in a falcon tube. (B) Collagen-cell solution is transferred into the mould and allowed to gel in the incubator at 37°C, 5% CO_2_, and 90% humidity. (C) The mould and gelled collagen-cell solution are raised from the substrate, submerged in media, and incubated for a further 2 days before testing. (D) The tissue is removed from the mould and transferred to the modified biaxial test machine for experimentation. The same procedure is used to prepare both biaxial (upper panels of B-D) and uniaxial (lower panels of B-D) tissues. (E) Key dimensions for the uniaxial (left) and biaxial (right) moulds. (F) Nominal loading strain, *ε*_*L*_, applied to the contracted collagen-cell gel (tissue) at the hangers for the first 3 seconds of the cyclic loading step.

### 2.2 Testing System & Experiments

Significant modifications are made to a biaxial tensile tester (Zwick/Roell, Ulm, Germany) in order to perform the cyclic experiments on tissues. In order to measure bi-directional force, two load cell transducers (AE801, Kronex, CA, USA) are wired to a custom-built circuit that provides the relevant excitation voltage and amplification of the output signal. Digitisation, calibration, and data acquisition of the output signal are performed using CompactRIO and LabVIEW software (National Instruments, TX, USA).

After 2 days of incubation, tissues are imaged before being transferring to the modified test machine. Due to the contractile effects of the embedded cells, gels shrink during the 2-day incubation period to form “tissues”. After releasing hangers from the mould the tissues immediately spring/contract inwards. After loading into the modified test machine, manual adjustments are made to return the tissue to its original “mould” geometry (Figure 1-D). It should be noted that the tissue remains submerged in growth media at all times. After loading into the modified test machine, the sample is allowed to stabilise for 30 minutes before experimentation. After an initial loading step to 20% nominal strain in the first second, sinusoidal cyclic deformation is applied to the tissue between 10-20% nominal strain at a frequency of 1 Hz for 2 hours (Figure 1-F), applied uni-directionally for uniaxial specimens and bi-directionally for biaxial specimens. After 1 hour, 5 μM cytochalasin-D (cytoD) is added to the surrounding growth media to disrupt contractile SFs in tissues. Uniaxial specimens are free to contract laterally at all times. During cyclic deformation, measured force is recorded at a rate of 40 Hz. Measured experimental forces *F*_*E*_ are normalised such that the *effective nominal stress* (ENS) is defined as 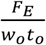, where the undeformed tissue cross-section width *w*_*o*_ and thickness *t*_*o*_ at the hangers are the same for both uniaxial and biaxial specimens (i.e. thickness *t*_*o*_ = 3*mm*; width *w*_*o*_ = 10*mm*). The reader is referred to the study of Nolan and McGarry (2016) for analysis and discussion of the nonuniformity of the stress state in biaxial specimens and the limitations of interpreting a normalised force as a measure of material stress. Measurements are subjected to statistical significance testing. Statistical analysis is performed using GraphPad (GraphPad, CA, USA) using a two-way analysis of variance (ANOVA). For all comparisons, statistical significance is declared if p < 0.05.

### 2.3 Immuno-fluorescence

Tissues are fixed by submerging them in 4% paraformaldehyde solution for 20 minutes. Fixed tissues are treated with 0.1% Triton solution for 5 minutes. FITC phalloidin and DAPI dilactate (Sigma–Aldrich Ireland Ltd., Arklow) are used to stain the actin filaments and nuclei, respectively. Washing 3 times with PBS is performed between each step. Samples are slightly agitated on a rotation plate during each step to assist penetration of reagents throughout the tissues. Samples are mounted in Prolong Gold Antifade Reagent (Molecular Probes, Life Technologies, NY, USA) to preserve staining quality and kept below 4°C before imaging. An Andor Revolution spinning disk confocal microscope (Yokogawa CSU-X1 Spinning disk unit & Olympus IX81 microscope) is used to image fluorescently stained samples. To investigate the SF distribution, representative fluorescent z-stack images are obtained at 5 regions of biaxial and uniaxial tissues before and after experiments. Two image channels are captured at each plane of focus, using laser excitation of 405 nm to visualise the nucleus and excitation of 495 nm to visualise the actin filaments. A step size of 2 m is used between each z-plane. Complete z-stack image sets are grouped into a single image using maximum intensity. To quantify the SF orientation distribution in the tissues, fluorescent images obtained at key areas of interest are analysed using the OrientationJ plugin for image processing software ImageJ. SFs are grouped by their angles to the nearest 1° increment. Probability density functions of the resulting orientation distribution are then generated.

## 3 Experimental Results

### 3.1 Collagen-Cell Gel Contraction

After gelling, the collagen-cell solution samples are incubated for 2 days before mechanical testing. During this 2-day period cells elongate and spread within the three-dimensional collagen matrix and endogenous force generated by SFs leads to gel contraction. In Figure 2 a quantitative analysis of the deformation of uniaxial and biaxial tissues is shown. In uniaxial tissues the width of specific sections along the length are measured (Figure 2-A). For each measurement a significant reduction from the baseline (day 0) width is observed (Figure 2-B). Mid-way between the opposing hangers a minimum width of WM = 4.3±0.3 mm is measured. This dramatic reduction in sample width correlates to a large negative nominal strain (−57%) in the lateral direction. The width gradually increases from the centre section to 4.5±0.3 mm at W3 (3 mm from the mid-way section), 5.3±0.4 mm at W2 (6 mm from the mid-way section), and 7.3±0.5 mm at W1 (9 mm from the mid-way section). On the hanger, contractile forces reduce the width of the sample to 8.6±0.5 mm (WH). The non-uniform contracted width along the length of the sample is due to a partial constraint in the lateral direction as a result of friction between the tissue and the hangers. In the case of the biaxial tissue, the tissue is unconstrained only in the corner regions between orthogonal hangers. Deformation of the biaxial tissue during static incubation is characterised by the change in the distance, LC, indicated in Figure 2-C. In the absence of contractility LC=1.707 mm. As shown in Figure 2-D, LC increases to 2.22±0.27 mm due to cell contractility.

**Figure 1:**
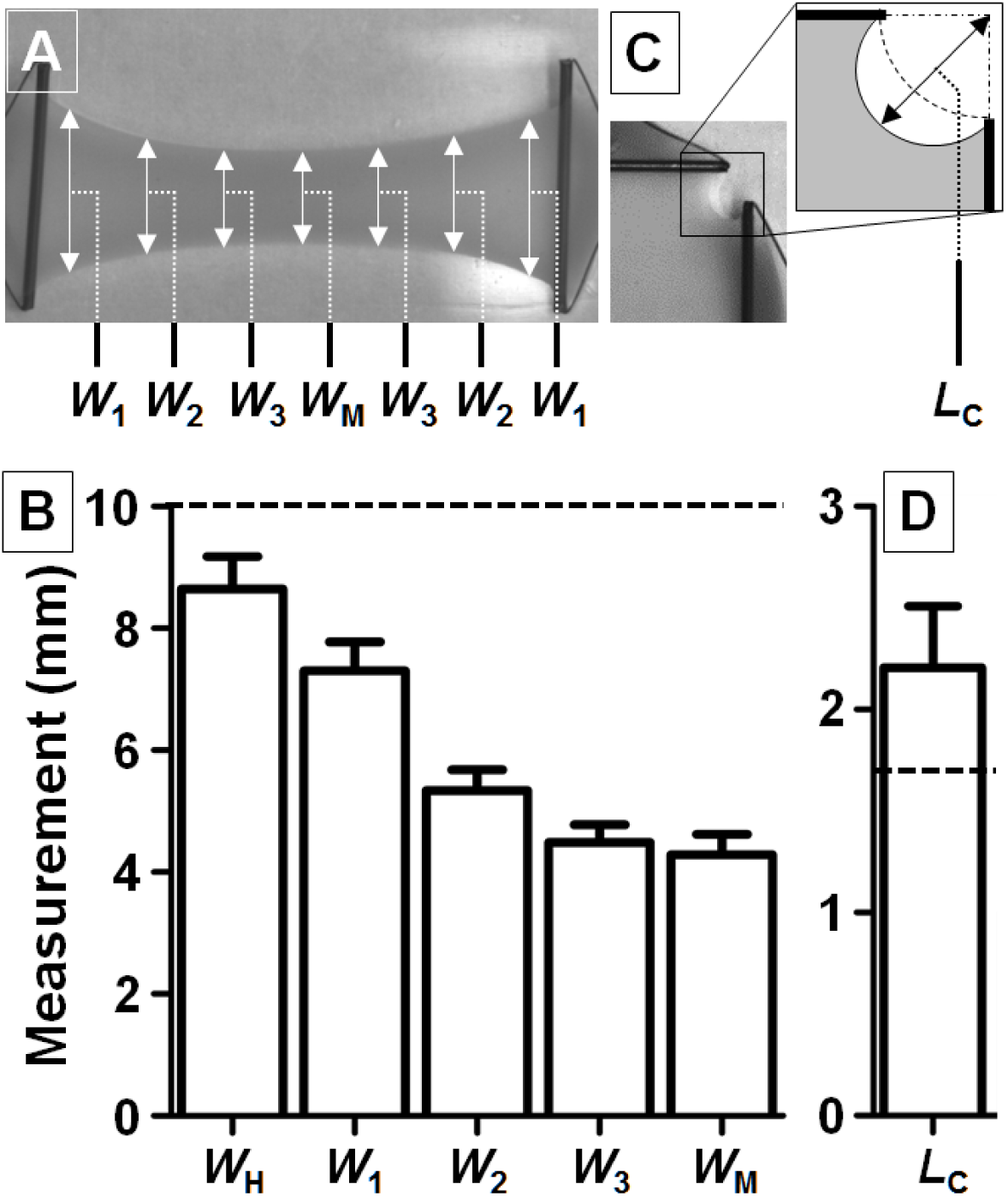
Measurements of contractility mediated deformation of biaxial and uniaxial specimens after 2 days of static incubation. (A) Contracted uniaxial tissue indicating regions at which width measurements are made. Distance between regions is 3 mm. Region W_1_ is 1 mm from the hangers. Note that width measurements are also made at the hangers, W_H_. (B) Plot of measurements (mean ± standard deviation, n=6) for the uniaxial tissues at each section. (C) Quadrant of a contracted biaxial tissue indicating the corner length, *L*_C_, measured after 2 days. (D) Measurement of the corner length, *L*_C_, of the biaxial tissue (mean ± standard deviation, n=6). In (B) and (D), baseline (day 0) measurements are indicated by dashed lines.

### 3.2 Maximum Nominal Stress Measurements

In Figure 3-A and 3-B the maximum ENS measured during cyclic deformation experiments are shown (full ENS-nominal strain loops are presented in Appendix A). The maximum force is recorded at the mid-point of each loading cycle, when tissue/hydrogels are at a maximum applied nominal strain of 20%. At the peak of the initial loading cycle (Figure 3-C), maximum ENS is 0.62±0.22 kPa for the uniaxial tissues, and 0.74±0.22 kPa for the biaxial tissues. For hydrogels (Figure 3-C), corresponding values of 0.24±0.3 kPa and 0.50±0.11 kPa are measured for the uniaxial and biaxial specimens, respectively. This demonstrates that the cells contribute a significant proportion of the measured force during the initial loading half-cycle in the case of both the uniaxial and biaxial tissues. Addition of cytoD to tissue samples reduces the measured ENS to 0.10±0.04 kPa and 0.14±0.02 kPa for uniaxial and biaxial samples, respectively (Figure 3-D). This further confirms the significant contribution of cell force in untreated contractile uniaxial and biaxial tissues.

**Figure 3:**
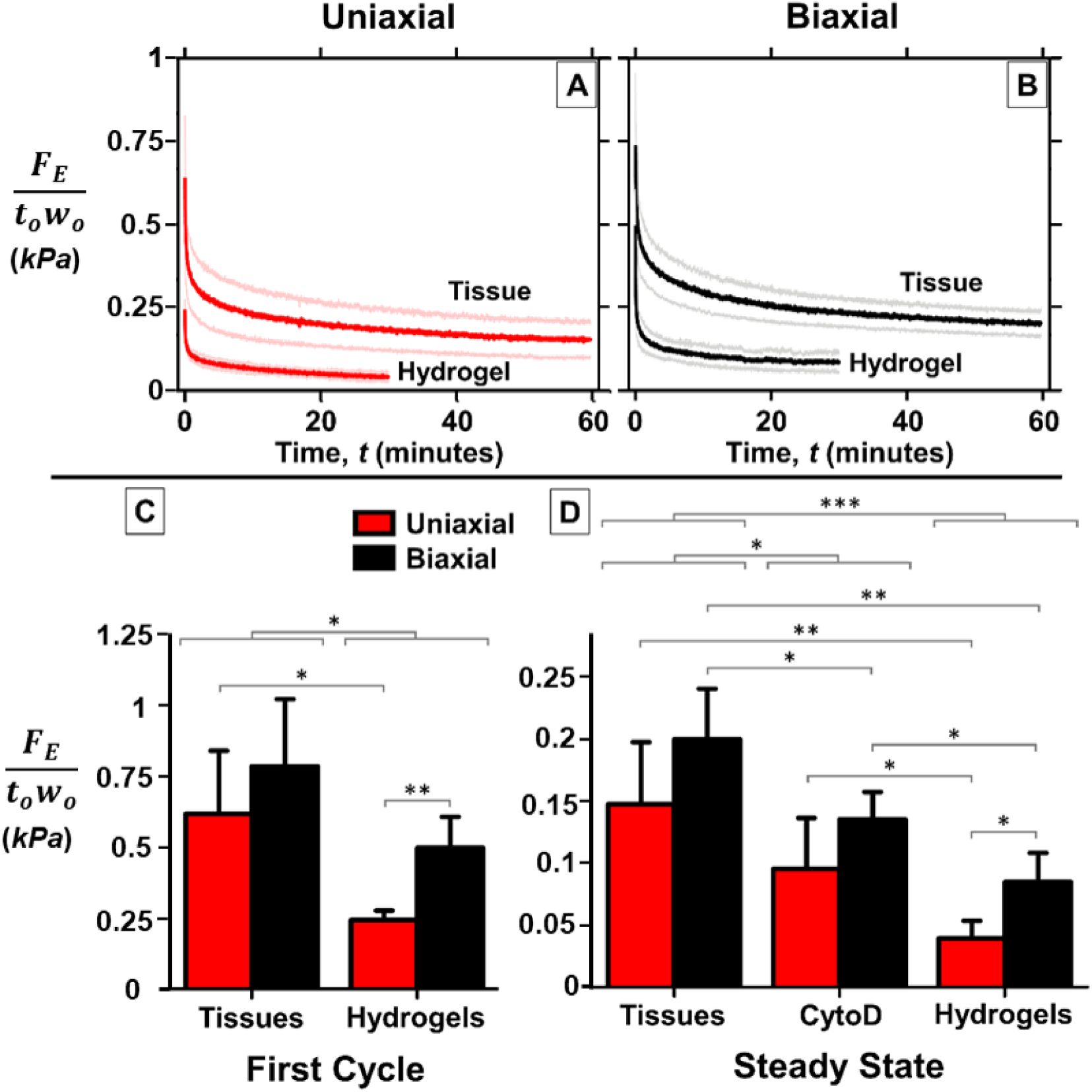
The maximum ENS measured from cyclic deformation experiments of: (A) uniaxial collagen tissues and hydrogels (Mean (red curve) ± standard deviation (pink curves), n=6) and (B) biaxial collagen tissues and hydrogels (Mean (black curve) ± standard deviation (grey curves), n=6). (C) The ENS measurement from uniaxial and biaxial deformation experiments at the end of the initial loading cycle for tissues and hydrogels. (D) ENS measurement after 60 minutes of cyclic loading of tissues, 60 minutes after the addition of cytoD, and after 30 minutes of cyclic loading of hydrogels. Note that maximum ENS is recorded when samples are at maximum stretch of 20% nominal strain. Statistical significance: *p<0.05, **p<0.005, ***p<0.0005..

A relaxation of maximum ENS to a near steady state value is observed for both uniaxial (Figure 3-A) and biaxial (Figure 3-B) hydrogels over the course of 30 minutes of cyclic loading. This demonstrates that the hydrogel behaves as a viscoelastic material.(Nam et al., 2016, Ban et al., 2018). A similar stress relaxation is also observed for the tissue, primarily due to the passive viscoelastic contribution of the hydrogel to the measured force. At steady state a maximum ENS of 0.15±0.05 kPa is measured for the uniaxial tissue, compared to a value of 0.20±0.04 kPa for the biaxial tissue (Figure 3-D). For hydrogels at steady state, significantly lower ENS values of 0.04±0.01 kPa and 0.08±0.02 kPa are measured for uniaxial and biaxial hydrogels, respectively. This again confirms the dominant contribution of cells to the steady state stress in both uniaxial and biaxial tissues. The change of ratio of biaxial to uniaxial steady state ENS from 2.08 for hydrogels to 1.36 for tissues again highlights the dominant role of cell contractility, and further underscores that the mechanical contribution of cells cannot be simplistically interpreted as an increase in the tissue stiffness.

### 3.3 Fluorescent Images

Fluorescent images of uniaxial and biaxial tissues are shown in Figure 4 and Figure 5, respectively. Five specific regions of interest are imaged in each tissue to investigate the SF distribution, both pre- and post-cyclic deformation experiments. Fluorescent images are generated by combining multiple z-stack images. 3D rendered image z-stacks reveal that cells are predominantly spread in the plane of the tissue (x-y) for both biaxial and uniaxial samples, with cells tending to align orthogonal to the out-of-plane direction. Furthermore, cells and their SFs become highly elongated throughout the uniaxial and the biaxial tissue. However, the alignment of cells and SFs is shown to be highly dependent on the applied constraint, as described below.

**Figure 4:**
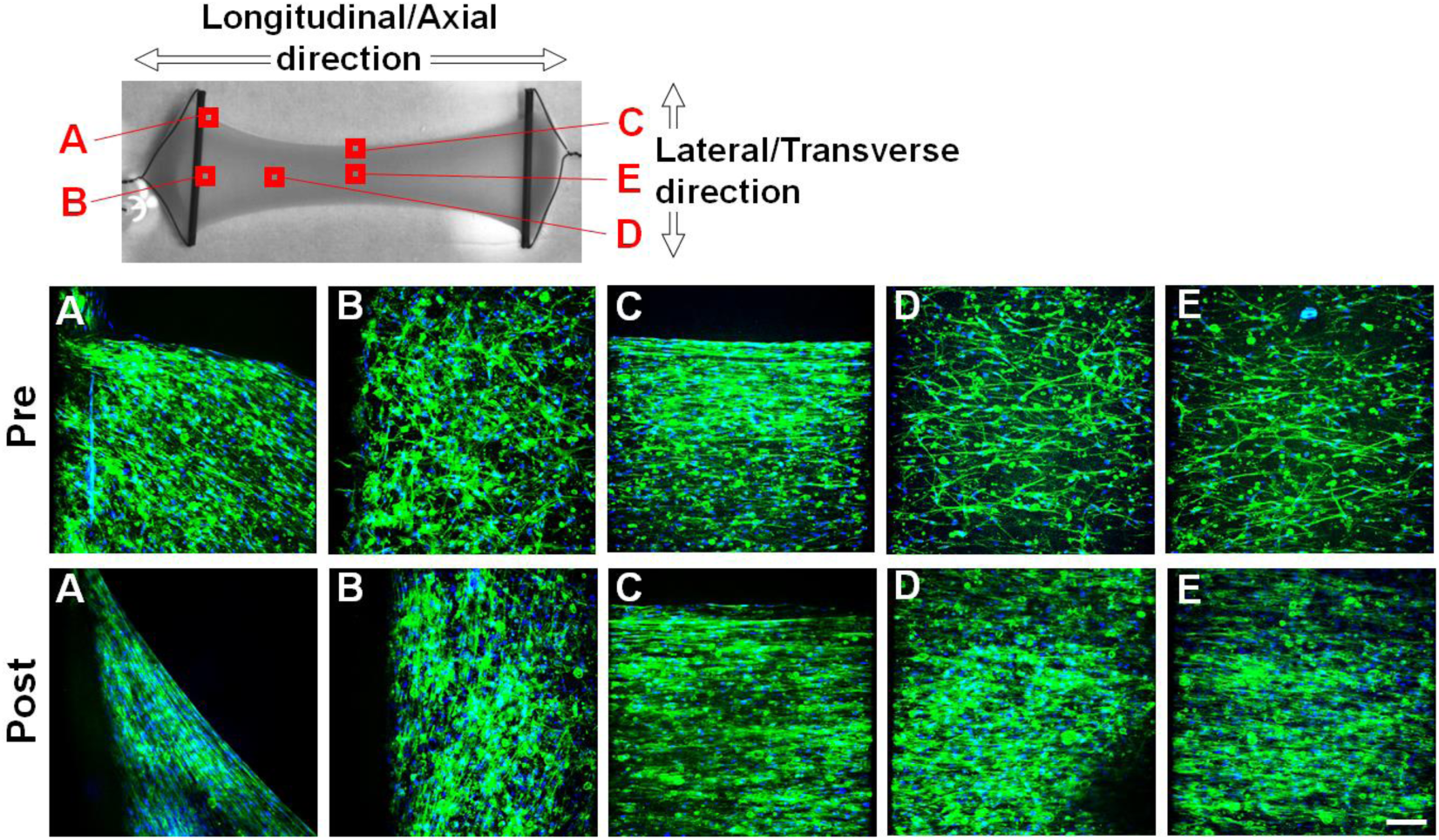
Fluorescent images of the uniaxial tissues. Actin is labelled with green and nuclei are blue. The maximum intensity of multiple z-stacks are combined to create images at the following regions: (A) the edge of the tissue-hanger interface region, (B) the middle of the tissue-hanger interface region, (C) the edge of the tissue between the two opposing hangers, (D) a region mid-way between the hanger and the centre of the tissue, and (E) the centre of the tissue. Scale bar = 250 μm.

**Figure 5.**
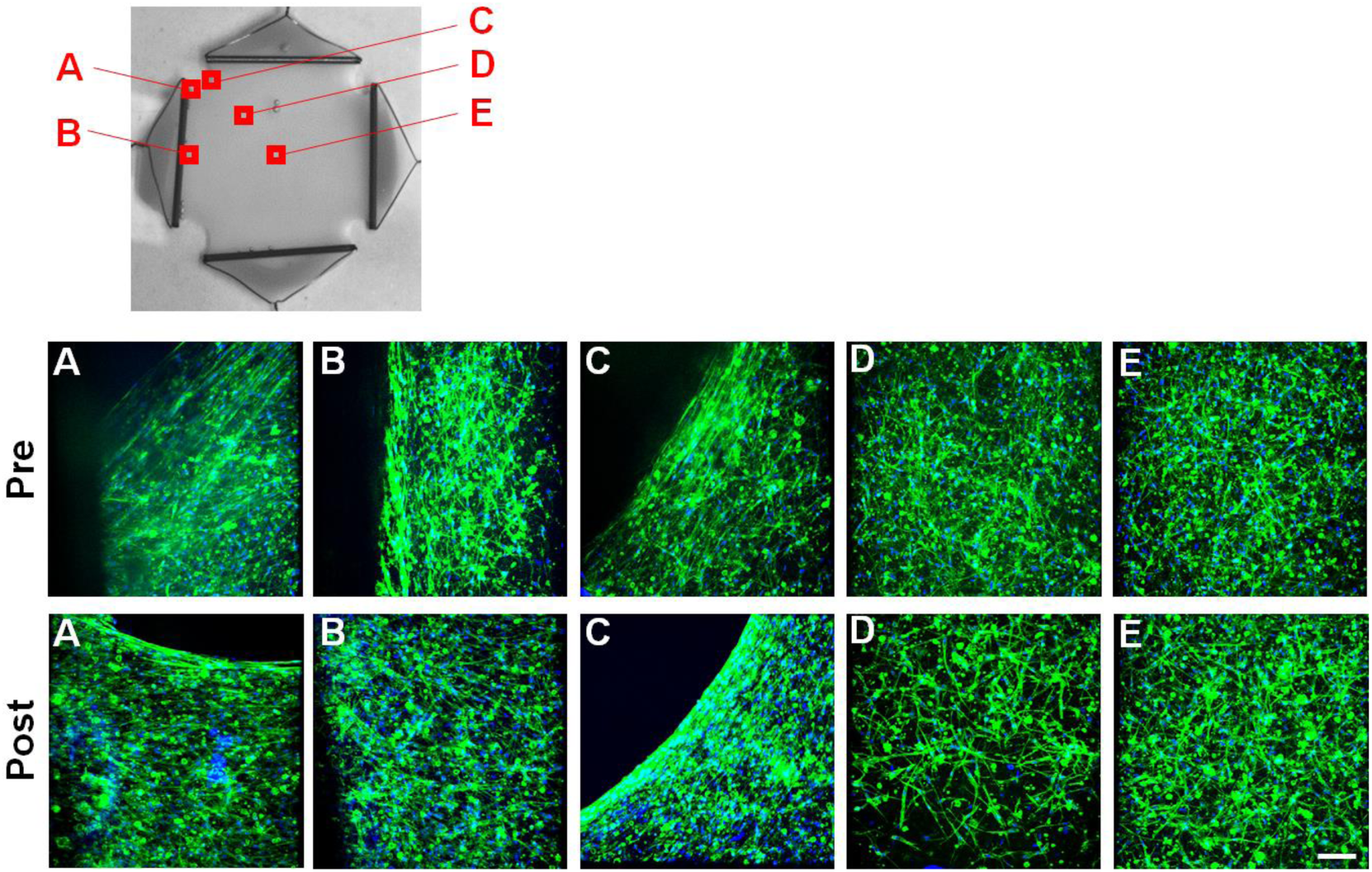
Fluorescent images of the biaxial tissues. Actin is labelled with green and nuclei are blue. The maximum intensity of multiple z-stacks are combined to create images at the following regions: (A) the edge of the hanger-tissue interaction region, (B) the middle of the hanger- tissue interaction region, (C) the edge of the tissue between two neighbouring hangers, (D) a region mid-way between the edge and the centre of the tissue, and (E) the centre of the tissue. Scale bar = 250 μm.

In the case of the uniaxial tissue (Figure 4), significant alignment of cells and their SFs in the longitudinal direction is observed, even before the application of cyclic stretching, as shown in Figure 4-C, D (pre). Following 1 hour of cyclic stretching, the alignment of SFs in the longitudinal direction is further enhanced (Figure 4-C, D (post)). As previously demonstrated in Figure 2, lateral contraction of the tissue is constrained near to the hangers, which has a significant influence on the local SF alignment. At region A (Figure 4-A), SFs predominantly align parallel to the free edge of the tissue, both pre- and post-cyclic loading. However, at region B, before cyclic loading is applied (Figure 4-B (pre)) SFs align in the lateral direction at the tissue-hanger interface, but become randomly aligned farther from the hanger. Application of cyclic loading (Figure 4-B (post)) further enhances the lateral alignment of SFs at the tissue-hanger interface.

In Figure 5-D and -E, it is clearly shown that cells/SFs are randomly oriented throughout the majority of the biaxial tissue, both pre- and post-biaxial cyclic loading. As shown in Figure 5-C highly localised cell/SF alignment is observed parallel to the free-edges at the corner regions of the tissue (Figure 5-C (pre)). The random alignment is restored a short distance from the free edge. Application of biaxial cyclic stretching further enhances alignment parallel to the corner free-edges. In region B (Figure 5-B (pre)), a localised alignment of cells parallel to the hanger is observed at the tissue-hanger interface. Interestingly, in contrast to the uniaxial tissue, application of biaxial cyclic stretching disrupts this localised alignment at the tissue-hanger interface. At the intersection between the hanger and the free-edge (region A, Figure 5-A) SFs are observed to align parallel to the free edge, rather than parallel to the hanger.

In Figure 6, a quantitative analysis of SF distribution in key regions of uniaxial and biaxial tissues is presented. At region A (Figure 6-A) in the uniaxial tissue, on the free-edge mid-way between the two opposing hangers, the SF distribution is highly peaked pre-cyclic stretching. Cyclic stretching slightly increases the peak of the distribution in the longitudinal direction. At region B (Figure 6-B) in the centre of the uniaxial tissue, the SF distribution is moderately peaked in the longitudinal direction pre-cyclic stretching. Cyclic stretching increases the peak of the distribution, indicating that a further SF realignment occurs during applied deformation. At the free edge in the corner region of the biaxial tissue a highly peaked SF distribution is observed pre-cyclic stretching (Figure 6-C). Application of cyclic stretching further increases the peak of the distribution parallel to the free edge. As shown in Figure 6-D, in the central region of the biaxial tissue an approximately uniform distribution of SFs across all directions is observed. Application of cyclic stretching has no influence on the SF distribution in this region.

**Figure 6.**
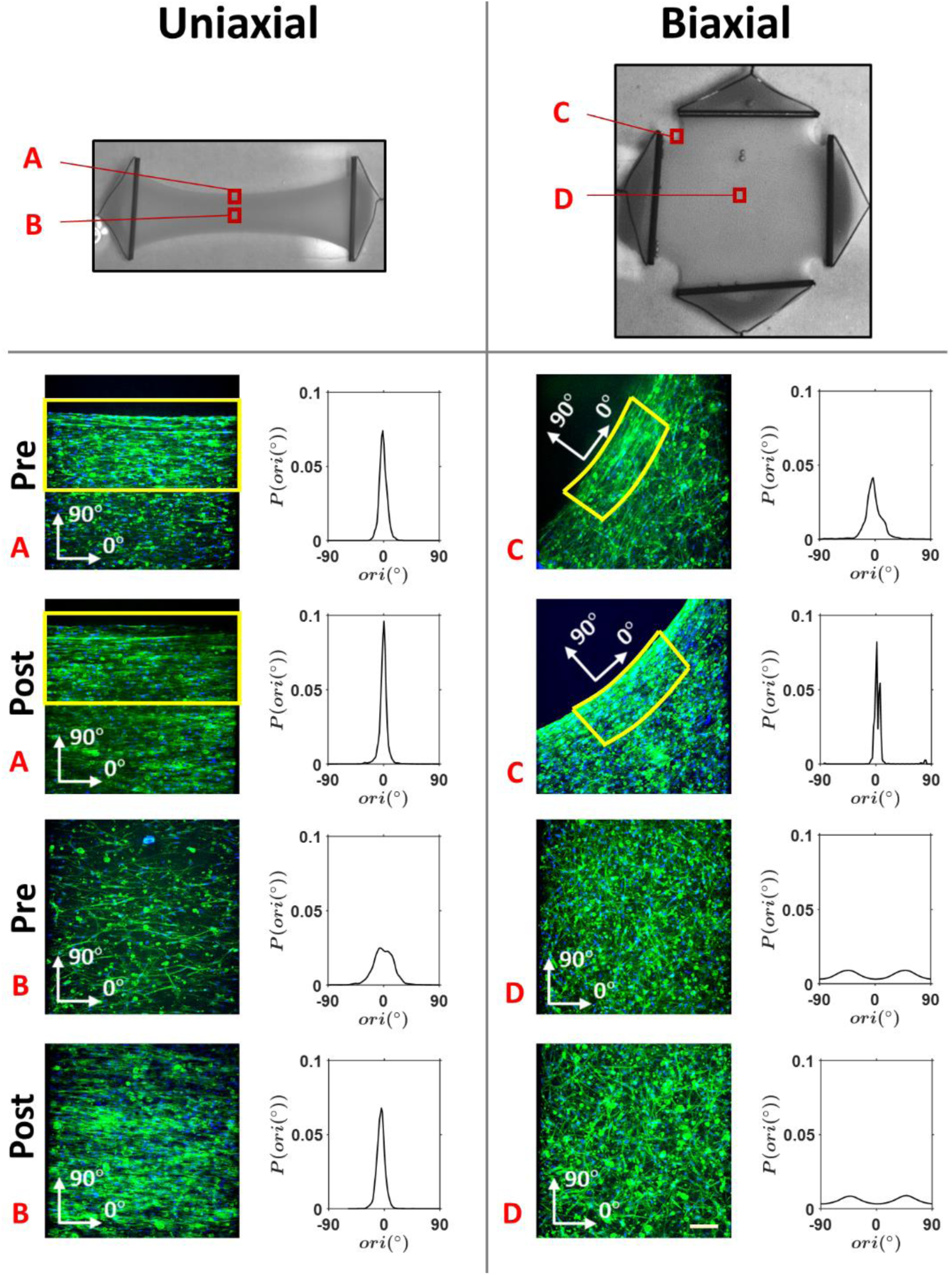
Fluorescent images and associated normalised SF orientation distribution at specific regions of uniaxial and biaxial tissues are shown. Actin is labelled with green and nuclei are blue. In the uniaxial specimens, normalised SF orientation distribution is measured from fluorescent images obtained pre- and post-experimentation at: (A) the edge of the tissue mid-way between the two opposing hangers, and (B) the absolute centre of the tissue. In the biaxial specimens, normalised SF orientation distribution is measured from fluorescent images obtained pre- and post-experimentation at: (C) the edge of the tissue between two neighbouring hangers, and (D) the centre of the tissue. In (C), 0° SFs are those that align tangentially to the edge of the specimen. In (A) and (C), only a limited region of interest considered (outlined in yellow). The regions of interest for uniaxial and biaxial specimens have a depth of 300 □m and 500 □m from the edge, respectively. Scale bar = 250 μm.

## 4 Computational Analysis

The experimental measurements presented above suggest that active cell contractility significantly contributes to the measured force for both biaxial and uniaxial tissues. Experiments also reveal significantly different patterns of cell alignment in uniaxial and biaxial tissue. However, the active stress generated by cells in uniaxial and biaxial tissues can only be parsed through a model framework that accounts for active cell contractility and alignment.

Finite element simulations (Abaqus, Dassault Systems, RI, USA) of the experiments are performed to investigate stress-fibre remodelling and collagen realignment in the tissue constructs. The novel tissue model entails the simulation of active cell contractility and alignment using the thermodynamically consistent active cell models of Vigliotti et al., (2015) and McEvoy et al. (2017) in parallel with a 3D anisotropic collagen gel. The collagen gel is modelled using an anisotropic hyperelastic framework, where the initially uniform distribution of collagen fibres can alter its distribution and alignment due to the contractile action of the cells (or due to an externally applied displacement). Such collagen alignment in turn results in a further alignment of cells, providing a thermodynamically motivated mechanistic representation of contact guidance.

Full details of the computational model are provided in Appendix B. In summary, we consider SF formation in *ϕ*_*n*_ discrete directions. The active stress tensor due to cell contractility is given as:

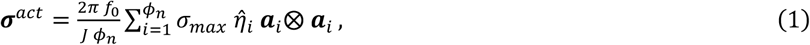

where 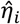 is the cross-sectional SF concentration in direction *i, σ*_*max*_ is the maximum isometric tension in a SF, *f*_0_ is the cell volume fraction in the tissue, *J* is the determinant of the deformation gradient ***F***, and ***a***_*i*_ = ***F a***_0*i*_, where ***a***_0,*i*_ is a unit vector indicating direction *i. ϕ*_*n*_ = 240 is found to result in a converged active stress tensor. The value of the cell volume fraction, *f*_0_, is calibrated to match the experimentally measured uniaxial tissue force. The calibrated value is expected to correspond to the experimentally applied cell densities (1×10^6^ cells per ml hydrogel, see Section 2.1). Predicted cell distributions in the tissue are characterised in terms of the cross-sectional concentration of sarcomeres in a given direction *i*, 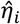, and also in terms of the total sarcomere formation in a given direction *i*, 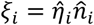, where 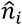 is the number of sarcomeres in series (see Appendix B). The mechanical behaviour of the collagen hydrogel is described by a hyperelastic model incorporating dispersed fibres, following the approach of Holzapfel et al. (2002). Here we assume an initially isotropic distribution of collagen fibres in an undeformed tissue. The passive anisotropic stress tensor is computed by numerically integrating the stress contribution of *ϕ*_*m*_ collagen fibres (Li et al., 2018; McEvoy et al., 2018), such that

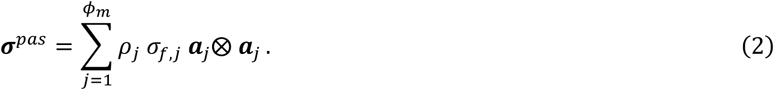

*ρ*_*j*_ is the collagen fibre concentration in direction *j*, and *σ*_*f,j*_ is the collagen fibre stress in direction *j*. The passive framework is completed by the addition of a compressible isotropic component, following the approach of Nolan and McGarry (2016a).

A finite element mesh of the undeformed collagen is constructed based on the mould geometries described in Section 2.1. The 2-day incubation period is simulated by applying zero displacement boundary conditions to the edges of the tissue that are constrained experimentally by the hangers. Following this step, displacement boundary conditions are applied to the post-incubation tissue configuration, as implemented in the biaxial and uniaxial experiments described in Section 2.2. Actively generated cell contractility and remodelling is simulated in parallel to collagen deformation and realignment. Computed tissue deformation and tissue forces are compared to experimental measurements. Simulations are also performed for passive collagen hydrogels by removing the active cell component of the model. Computed results are again compared to the experimental data for cell-free hydrogels.

### Computational Results

As shown in Figure 7-A, the passive anisotropic hyperelastic hydrogel is calibrated using the uniaxial model so the experimentally measured ENS (0.04 kPa) is exactly computed. Using these calibrated passive hydrogel properties, the predicted ENS for the biaxial hydrogel model (0.087 kPa) is close to the experimentally measured value (0.08±0.02 kPa), suggesting that the passive anisotropic hyperelastic model accurately captures the degree of collagen fibre alignment during uniaxial stretching, as shown in Figure 7-B. The active cell contractility model is added to the passive anisotropic hyperelastic model and the cell volume fraction of *f*_0_ = 0.008 is calibrated so the experimentally measured ENS (0.15 kPa) for a contractile cell seeded tissue is exactly computed, as shown in Figure 7-C. This value of *f*_0_ is within the expected range based on the experimental cell density of 1×10^6^ cells per ml hydrogel. The calibrated active-passive model predicts a biaxial ENS of 0.22 kPa, which is close to the experimentally measured value (0.20±0.02 kPa), as shown in Figure 7-C. demonstrating the predictive capability of the model in terms of cell contractility and sarcomere alignment. The addition of the active cell component to the hydrogel model significantly decreases the ratio of biaxial to uniaxial force (from 2.08 (passive hydrogel) to 1.45 (active cells + hydrogel)), as observed experimentally. The model reveals that this ratio decreases due to the following reasons: (i) As shown in Figure 7-D, high levels of cell sarcomere alignment in the loading direction in the uniaxial tissue results in a higher total active cell Cauchy stress component in the loading direction at the centre of the tissue 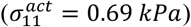. In comparison, sarcomere formation in the biaxial tissue is uniformly distributed across all orientations, resulting in a lower active cell Cauchy stress component in the loading directions at the centre of the tissue 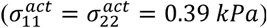, despite the fact that computed total sarcomere formation is slightly higher (∼3%) in the biaxial tissue. (ii) In uniaxial tissues, high levels of collagen alignment occurs due to cell contractility. Such pronounced collagen alignment does not occur in hydrogels subjected to uniaxial loading. This cell induced collagen alignment increases the apparent passive stiffness of the uniaxial tissue in the loading direction. In summary, the model reveals that the ratio of biaxial to uniaxial force is significantly higher for a passive hydrogel than for a contractile tissue due to alignment of sarcomeres in uniaxial loading conditions, in addition to cell induced collagen alignment.

**Figure 7.**
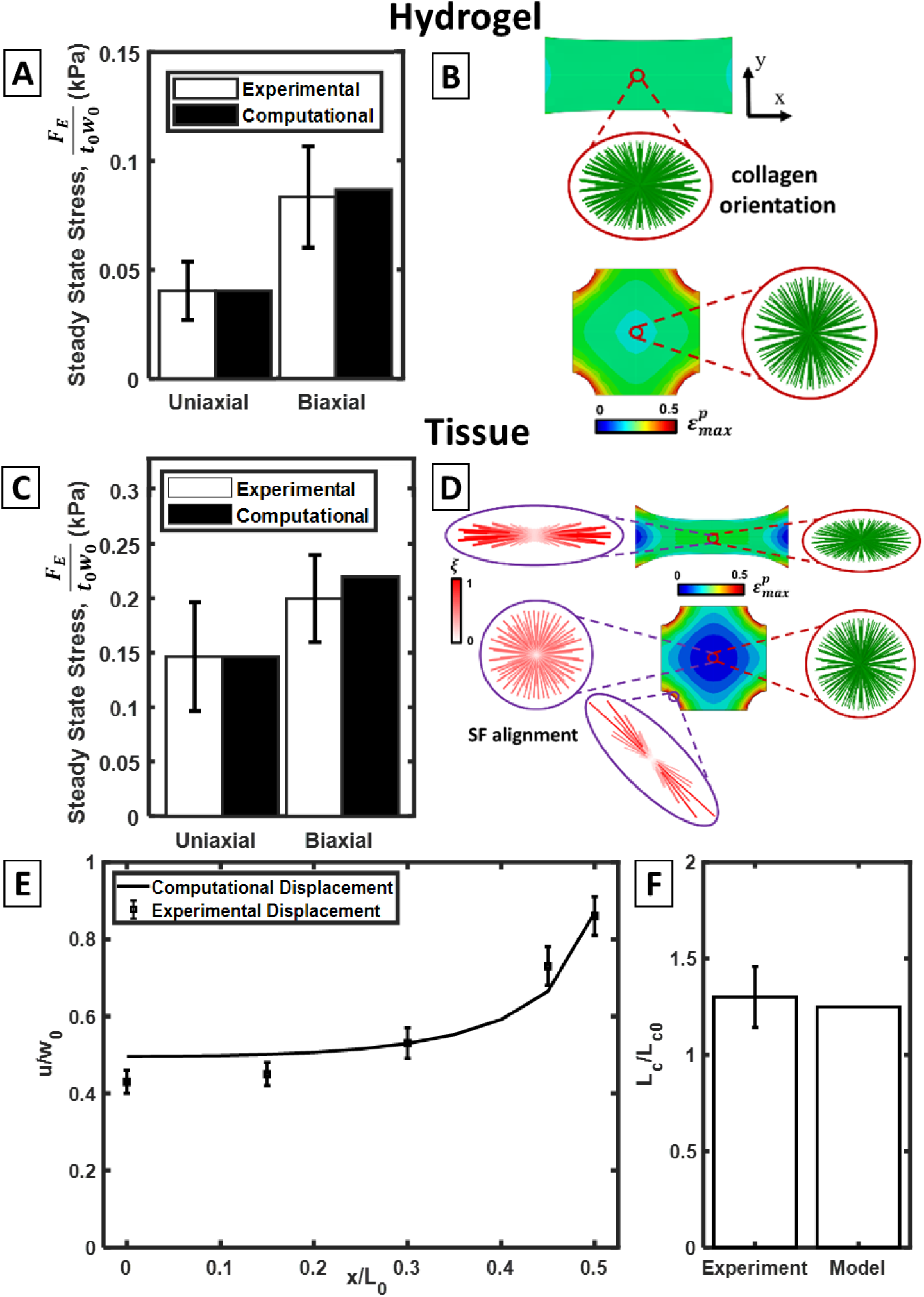
(A) Experimental and computational steady state ENS for hydrogels. (B) Collagen alignment (green) in stretched (*ε*_*L*_= 20%) uniaxial and biaxial hydrogels. Contour plot shows distributions of max principal strain throughout the hydrogel; (C) Experimental and computational steady state ENS for tissues; (D) Total sarcomere formation 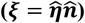 in a given direction (red) and collagen alignment (green) in stretched (*ε*_*L*_= 20%) uniaxial and biaxial tissues. Both sarcomere line length and colour intensity scale with *ξ*, while sarcomere line thickness scales with cross-sectional concentration of sarcomeres, 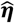 Contour plots show distributions of max principal strain throughout the tissue; (E) Experimental and computational unloaded steady state normalised contracted width in uniaxial specimens; (F) Experimental and computational steady state unloaded normalised contracted corner length in biaxial specimens.

The model is further validated by the accurate prediction of cell alignment along stress free boundaries at the corners of biaxial gels (Figure 7-D) in contrast to unaligned cells throughout the majority of the central region of biaxial tissues. This prediction is in strong agreement with experimental data presented in Figure 6, suggesting that the model can be used to predict the influence of heterogeneous multiaxial stress states on cell alignment throughout a tissue. As a final validation of the model, Figure 7-E demonstrates that the model accurately predicts the deformation of the uniaxial tissue due to cell contractility under conditions of static incubation. The deformation of the stress-free boundaries of the biaxial tissues are also accurately predicted (Figure 7-F). This further suggests that the model captures the key physics of the interaction between the anisotropic passive hyperelastic hydrogel model and the active model of cell contractility and remodelling. Additional analysis of the transient behaviour of the hydrogel and tissue are presented in Appendix C.

## 5 Discussion

In this investigation we develop a novel experimental methodology that allows for the observation of cell alignment, while characterizing cell contractility through measurement of active force in both uniaxial and biaxial tissues under dynamic loading. We also implement a novel modelling framework to parse the relationship between active stress generation and cell alignment observed in our experiments. Our characterisation of the coupling between cell contractility and alignment in biaxial and uniaxial dynamically loaded tissues advances on the recent investigation of Chen et al. (2018), which focuses on the influence of a biaxial constraint on alignment without characterisation of active contractility. The following key findings are uncovered:

- Addition of contractile cells to collagen hydrogels dramatically increases the measured forces in uniaxial and biaxial constructs under dynamic loading. This increase in measured force is due to active cell contractility, as is evident from the reduction in measured force after inhibition of actin polymerization with cytochalasin-D.
- Cells and their SFs are highly aligned in uniaxial tissues but are uniformly distributed in biaxial tissues, demonstrating the importance of tissue constraint on cell alignment.
- A novel active modelling framework reveals that while similar levels of SF formation occurs in both uniaxial and biaxial tissues, the uniaxial case results in a higher active cell stress due to three factors:
  i. Cells and SFs are highly aligned in uniaxial tissues, so that most actively generated contractile force contributes to the stress component in the loading direction.
  ii. The uniaxial gel becomes significantly compacted in the unconstrained direction. This increases the cell density across a plane orthogonal to the loading direction, in addition to the obvious effect of increasing the true stress.
  iii. Our model also accounts for the realignment of collagen due to cell contractility and mechanical loading. Cell contractility results in significant alignment of collagen in uniaxial tissues, resulting in a higher effective stiffness in this direction. This demonstrates the key interaction between active cell behaviour and passive hydrogel behaviour in determining the biomechanical tissue response.

By providing a quantitative link between constraint, cell alignment and actively generated cell contractile stress, the current study provides an important addition to the findings of Chen et al (2018), in which it is demonstrated that constraint is the predominant influence on cell alignment. The current study provides further evidence of this finding. Furthermore, through the decrease in force during uniaxial and biaxial dynamic loading with inhibition of actin polymerization through cytochalasin-D, the current study demonstrates that cells contributes significantly to the tissue force under both uniaxial and biaxial constraint.

Inhibition of actin with cytochalasin-D disrupts both active contractility (e.g. stress fibres) and passive forces (e.g. cytoskeleton). This supposes a limitation of the current study, as the contribution of active contractility cannot be quantified. However, one finding suggest that active contractility may be in place. The difference in the ratio of biaxial to uniaxial force for hydrogels is significatively different (2.04) to the one for tissues (1.20). If the cell contribution was simply a passive stiffening of the tissue then it seems reasonable that this ratio would not change, However, the difference in this ratio may be caused by the random alignment of cells observed in biaxial stretched samples, even if all the contribution would be passive.

As an alternative to cytochalasin-D, blebbistatin can also be used to parse active contractility of cells (Kovacs et al., 2004). However, blebbistatin has some limitations. Several studies have reported that the reduction in measured cell force due to blebbistatin treatment is not significantly different to the reduction in cell force due to cytochalasin-D treatment (Watanabe-Nakayama et al., 2011, Kraning-Rush et al., 2011, Sayyad et al., 2015). It is not therefore clear that using blebbistatin instead of cytoD will have any effect on the results of the current study. Cytochalasin-D is an inhibitor of actin polymerisation Therefore, in addition to removal of the active generated cell force, cytoD will also affect the passive contributions of actin cytoskeleton. However, several studies report that both cytoD and blebbistatin alter the morphology of the cells (Kraning-Rush et al., 2011, Sheets et al., 2013, Chen et al., 2010). Therefore, the passive strain energy stored in the cells consequently in the surrounding ECM will be altered by both cytoD and blebbistatin. Therefore, in addition to the intended effect of inhibiting active contractility, a secondary effect of both inhibitors will be a slight alteration of the passive cell/tissue forces.

Additionally, a detailed consideration of the molecular mechanism of blebbistatin inhibition also suggests that passive stiffness, in addition to active stress generation, tractions will be altered at a sarcomere level. Blebbistatin selectively inhibits the phosphate release from the myosin–ADP–Pi complex (Rahman et al., 2018), causing accumulation of cross-bridges and trapping the myosin motor in the pre-power stroke state, resulting in a product complex with some affinity for actin (albeit a lower affinity than in untreated cross-bridges (Sirigu et al., 2016). Therefore, although blebbistatin removes active traction generation, the passive stiffness of the actin cytoskeleton will also be altered due to the formation of non-stroking “passive” cross-bridges between actin and myosin. Such non-stroking “passive” cross-bridges will alter the passive structural stiffness of sarcomeres, in comparison to untreated cells.

Our active modelling framework accurately predicts our experimental trends and suggests that a slightly higher (3%) total SF formation occurs at the centre of a biaxial tissue compared to the uniaxial tissue. However, high alignment of SFs and lateral compaction in the case of the uniaxial tissue results in a significantly higher (75%) actively generated cell contractile stress, compared to the biaxial tissue. Previous studies (Thavandiran et al., 2013) reveal the influence of uniaxial and biaxial constraint on the alignment and SF formation in engineered cardiac micro-tissues. However, these investigations did not measure or analyse the resultant contractile force as a function of stress biaxiality and constraint. The current study provides new information on the link between tissue constraint, cell alignment, and actively generated contractile stress/force.

In the current study a very similar distribution of SFs is observed in tissues before and after the application of cyclic loading. In fact, we demonstrate that application of cyclic stretching results in a higher alignment of cells in uniaxial tissues, and an unchanged alignment in biaxial tissues. This demonstrates that the “strain-avoidance” hypothesis for cell alignment does not explain the influence of applied stretching on cell alignment in 3D scaffolds. Rather, our study suggests that tissue compaction due to non-biaxial constraints is the predominant influence on cell alignment. This explanation is also supported by the experiments of Chen et al. (2018). In fact, our active modelling framework suggests that cell contractility not only results in lateral compaction of a uniaxial gel, but also results in alignment of the collagen scaffold in the constrained direction. This further reduces the effective scaffold stiffness in the lateral direction, resulting in further cell alignment. Therefore, our model suggests that the commonly reported “contact guidance” hypothesis in 3D scaffolds is merely a demonstration of the influence of constraint on the coupling between cell and collagen alignment. This result is also supported by the experimental measurement of cell and collagen alignment in the study of patterns Rubbens et al. (2009). Furthermore, the highly aligned cells along the corner regions of our biaxial scaffold provide a further demonstration of this phenomenon. The alignment of cells in the direction of loading in uniaxially constrained 3D scaffolds has been widely reported in the literature (Foolen et al., 2012; Gauvin et al., 2011; Nieponice et al., 2007b; Wille et al., 2006; Zhao et al., 2013). Additionally, the reported experimental trend that cells align perpendicular to the direction of uniaxial stretching when seeded on 2D silicone substrates (Barron et al., 2007; Kaunas et al., 2005; Neidlinger-Wilke et al., 2001; Wang et al., 2001) can be explained in part by the observation that such 2D substrates are sufficiently thick and stiff that they cannot be compacted in the lateral directions by active cell contractility.

The current study presents a bespoke experimental approach to accurately measure tissue forces during uniaxial and biaxial dynamic loading and to relate measured force to cell alignment. Previous studies that report tissue forces during dynamic loading have largely been limited to uniaxial deformation. Zhao et al., (2013) fabricated contractile uniaxial collagen tissues on two opposing deformable microposts. By magnetically displacing a bead attached to one micropost, cyclic loading was applied to the tissues while the force was estimated by measuring the deflection of the opposing micropost. In the study of Wakatsuki et al., (2000), a system whereby tissue rings were fabricated for loading into a custom stretching rig is described. The force generated by the tissues rings during static contraction and during cyclic stretch was measured, in a similar fashion to the system described in the current study. In the studies of Zhao et al. (2013) and Wakatsuki et al., (2000), the mechanical contribution of cells during tissue stretch was quantified by comparing the force response of untreated tissues to tissues treated to disrupt the actin cytoskeleton. Furthermore, a similar technique is implemented to parse the response of the actin cytoskeleton in later studies (Wakatsuki et al., 2001; Wille et al., 2006). Again, it should be noted that these aforementioned studies are confined to uniaxial loading. Wagenseil et al. (2004) developed a system which investigated the anisotropic mechanical behaviour of collagen tissue vessels using a “pressure-diameter, force-length” test system. Vessels were either inflated to set diameters or axially stretched while the internal pressure, axial force, external diameter, and overall length were measured. It should be noted that in Wagenseil et al. (2004), tissue vessels were fabricated on rigid mandrels and, therefore, could only be engineered to exhibit circumferential or axial cell alignment before testing. Furthermore, the application of internal pressure and axial deformation was mutually exclusive. Measurement of force generated by biaxial tissues has generally been limited to a single applied monotonic stretch (e.g. Lee et al., (2008); Thomopoulos et al., (2007)), rather than a more physiologically relevant dynamic biaxial force. The bespoke experimental system developed in the current study facilitates: (i) performing testing on a broad range of tissue constructs with different engineered parameters, such as, collagen density and cell seeding density; (ii) the application of cyclic loading regimes to tissues; (iii) independent controlling of deformation in both stretching directions during biaxial testing; (iv) independent measurement of force in both stretching directions during biaxial testing; and (v) conducting immuno-fluorescent investigations of tissues before and after testing.

Previous experimental-computational studies have parsed the active and passive mechanical contributions of chondrocytes (Dowling et al., 2012, Dowling et al., 2013), endothelial cells (Reynolds et al., 2014), smooth muscle cells (McGarry et al., 2009), and osteoblasts (Reynolds et al., 2015; Weafer et al., 2013,15). The current study suggests that CMs exhibit high levels of active contractility, with cell alignment strongly influenced by multiaxial constraint.

Chen et al. (2018) quantify cell alignment in collagen constructs for uniaxial and biaxial loading. Thavandiran et al. (2013) seeded human embryonic stem cell differentiated CM in statically constrained uniaxial and biaxial collagen constructs. Cell behaviour is assessed by immunostaining for sarcomeric α-actinin, which is observed to be expressed in in the region of the stress-free boundaries in biaxial tissues, and expressed throughout uniaxial tissues. A comprehensive body of work by Sacks and co-workers investigates equi-biaxial dynamic loading and uniaxial strain (strip biaxial) dynamic loading of native tissues (Geest et al., 2005, Grashow et al., 2006, Liao et al., 2005) and tissue-engineered constructs (Gilbert et al., 2006, Courtney et al., 2006, Stella et al., 2008). Detailed analysis of observables such as ECM fibre orientation and nucleus aspect ratio are reported. The influence of pre-alignment of ECM fibres using electrospinning is also considered. The current study both compliments, and advances upon, these previous studies by measuring active cell and passive cell/tissue forces, in addition to cell alignment, in response to uniaxial and biaxial loading. Additionally, the implementation of a formulation for active cell contractility with a formulation for anisotropic collagen fibre reorientation within finite element simulations of our experiments allows for the computation of active and passive stress distributions within the tissue based on our experimental measurement of active and passive tissue forces. A study by Zhang et al. (2017) presents both active and passive force measurements for uniaxial CM-collagen constructs, in addition to alignment. The additional analysis of biaxial tissues in our study highlights the importance of multiaxial constraint on active force generation and CM alignment. Furthermore, following static tissue incubation, we observe short term (1 hour) changes in cell alignment in response to cyclic loading throughout uniaxial constructs. In biaxial constructs we observe such short term changes in cell alignment in the region of the stress free boundaries (where the local stress state is acts uniaxially), but we do not observe alignment changes in the central region of biaxial constructs (where the stress state is biaxial). Zhang et al. (2017) consider dynamic incubation, followed by quasi-static loading to characterise the properties of the tissue construct. A future study should combine the dynamic incubation protocols of Zhang et al. with the biaxial cyclic testing protocols presented in the current study to further explore the role of dynamic multi-axial stress states in the maturation and active contractility of engineered cardiac tissue. While the current focuses on the influence of multi-axial constraint and dynamic loading on alignment and contractility of immature cardiomyocytes, a follow-on investigation will consider higher cell densities within such multiaxially constrained tissues, in addition to longer term (days/weeks) application of dynamic multiaxial loading in an effort to accelerate the maturation of cardiomyocytes and enhance myotube and sarcomere formation.

Future studies will use the bespoke experimental rig developed here to investigate the influence of collagen density and anisotropy on cell alignment and active contractile force generation. In a study by Foolen et al. (2012), tissues that were originally biaxially constrained were subjected to a uniaxial cyclic stretch. If collagen density was “high”, the random SF distribution observed before cyclic loading was maintained in cells encapsulated in the core of the tissues. However, in tissues with a “low” collagen density, SFs exhibited stretch avoidance in the absence of a lateral constraint. Again, the bespoke experimental technique developed in the current study will allow for the measurement of tissue force as a function of collagen density. Furthermore, electrospinning techniques (Baker et al., 2008; Li et al., 2006) will also be used to directly manipulate collagen fibrils orientation to investigate the effect of contact guidance on cell alignment in tissue constructs.

In conclusion, our novel experimental-computational approach provides new insight into the relationship between tissue constraint, cell alignment, and active contractility. Experiments reveal that stress uniaxiality results in highly aligned SFs and cells throughout the tissue, whereas stress biaxiality results in randomly aligned SF distributions. In the case of both uniaxially- and biaxially-constrained tissues, cells and SFs are highly polarised and generate significant active contractile force. However, cells are highly aligned in uniaxial tissues but are uniformly distributed in biaxial tissues, demonstrating the importance of tissue constraint on cell alignment. A novel active modelling framework reveals that while similar levels of SF formation occurs in both uniaxial and biaxial tissues, the high alignment of cells and lateral compaction of the tissue in the uniaxial case results in a higher active cell stress, compared to the biaxial tissue.

## Acknowledgements

Funding was provided by Science Foundation Ireland (grants 18/ERCD/5481 and SFI/IP/1723). The authors would like to acknowledge the Irish Centre for High-End Computing (ICHEC) for provision of computational facilities and support. The authors thank Dr. Dimitrios Zeugolis for providing RTT collagen.

## Appendix A: Experimental Nominal Stress-Nominal Strain Loops

In Figure A1 the ENS measured during the cyclic deformation experiments are shown, with red (left) and black (right) plots representing mean values with standard deviations. The maximum ENS is recorded at the mid-point of each loading cycle, when tissue/hydrogels are at a maximum applied nominal strain of 20%. Individual loops for each experiment are also superimposed. In all cases a monotonic increase in stress is observed during loading and a monotonic decrease is observed during unloading. At the peak of the initial loading cycle (Figure A1-A), the mean ENS at peak applied strain is 0.62±0.22 kPa for the uniaxial tissues, and 0.74±0.22 kPa for the biaxial tissues. At steady state a mean ENS of 0.15±0.05 kPa at peak applied strain is measured for the uniaxial tissue, compared to a value of 0.20±0.04 kPa for the biaxial tissue (Figure A1-B). For cytoD treated tissues at steady state, significantly lower mean ENSs of 0.09±0.04 kPa and 0.13±0.02 kPa are measured for the uniaxial and biaxial specimens at peak applied strain, respectively (Figure A1-C).

**Figure A1.**
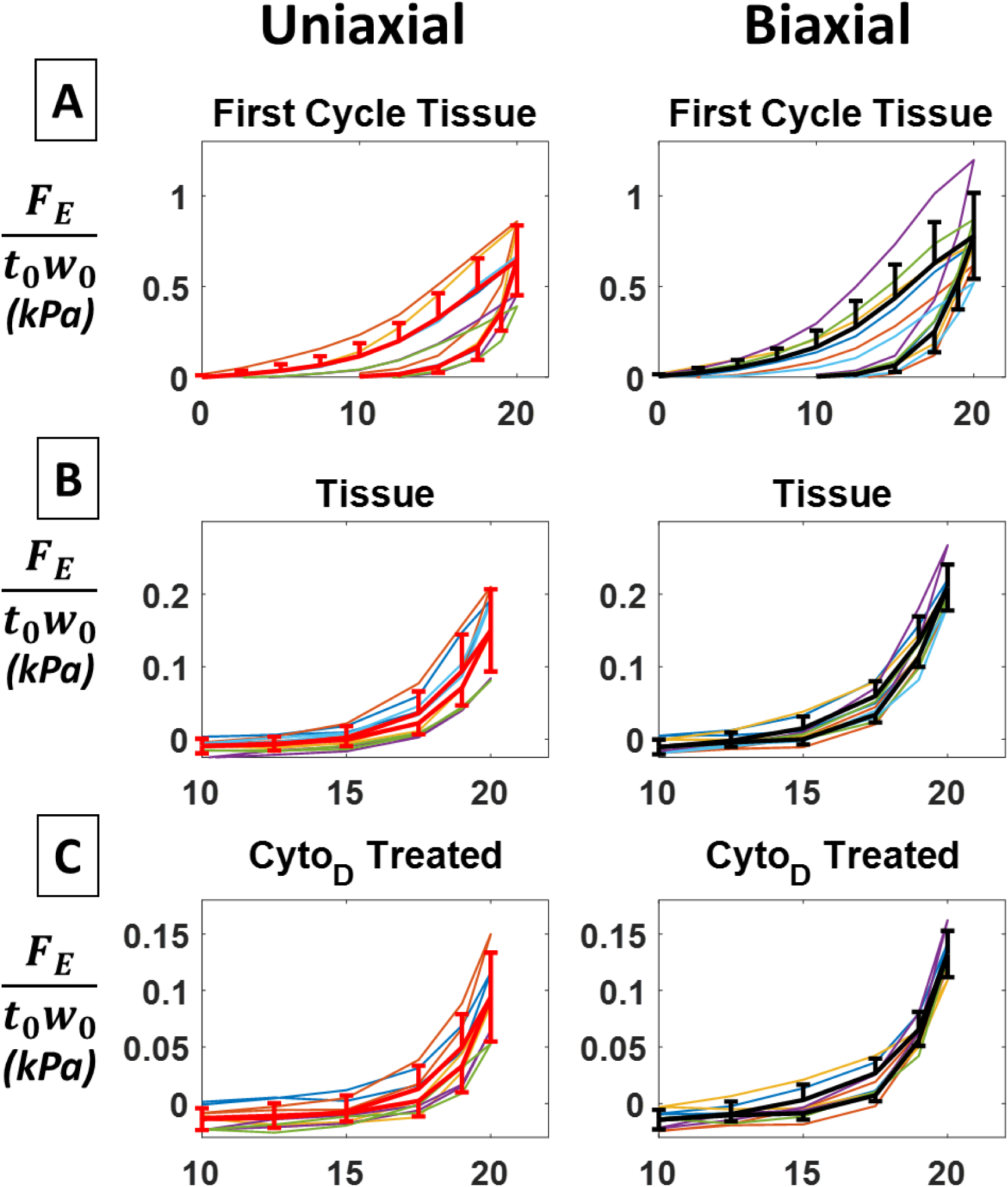
ENS results for uniaxial (left), and biaxial (right) cyclically loaded specimens, coloured line plots represent experimental data, with red (left) and black (right) line plots representing mean values with standard deviations. Individual loops of each experiment are also superimposed. (A) Tissue, during the first loading step; (B) Tissue during final steady state loading step; (C) CytoD treated specimens during the final steady state loading step.

## Appendix B: Experimental Nominal Stress-Nominal Strain Loops

### Cell model

The cellular cytoskeleton is composed of actin-myosin stress-fibres (SFs), which actively generate tension through cross-bridge cycling between the actin and myosin filaments. The thermodynamically consistent model from Vigliotti *et al*. (2015) captures key features of SF dynamics, including (i) the kinetics of SF formation and dissociation as motivated by thermodynamic considerations, (ii) the stress, strain, and strain-rate dependence of SF remodelling, and (iii) global conservation of the cytoskeletal proteins. Here we implement a steady-state form of this continuum model in a 3D finite element setting, following the derivations of McEvoy et al., (2017).

A representative volume element (RVE) in the undeformed state is defined as a sphere of radius *n*^*R*^*l*_0_/2. SFs emanate from the center of this sphere, each comprised of *n*^*R*^ axial sarcomeres (of length *l*_0_) in their initial ground state. In 3D, SFs can form in a large number of initially uniformly distributed directions *ϕ*_*n*_ (*ϕ*_*n*_ = 240 is found to provide a converged solution). At steady state, we consider that the (normalized) number of actin-myosin sarcomeres within a SF (in direction *i*) in the RVE is given by:

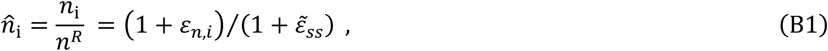

where *ε*_*n,i*_ is the nominal strain in the direction *i*. When a SF is extended, sarcomeres are added, with the effect that the internal strain in the SF is reduced until a steady state value 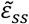 is achieved. 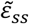 is given by the positive root of the relation:

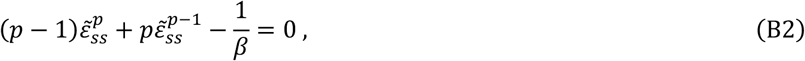

where *β* and *p* are non-dimensional constants that govern the internal energy *ψ* of *n*^*R*^ sarcomeres within a SF. Conversely, when a SF shortens, sarcomeres are removed. In both cases, the internal fibre steady state strain 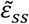 is fixed and in general different from the axial material strain in the direction of the fibre, *ε*_*n,i*_.

We assume the cells within the tissue construct are in the interphase period, when the cell is in a homeostatic state (i.e. the concentration of all proteins within the cell is constant) (Weiss, 1996). Therefore, in the finite element framework of this study a global conservation of the total number of SF proteins within the cell is enforced. Cytoskeletal proteins are considered to exist in two states: a bound state and an unbound state. The bound proteins make up the sarcomeres of the SFs within the RVE and thus are not mobile. The unbound proteins are mobile and can diffuse throughout the cell cytoplasm. The global conservation of cytoskeletal proteins may be expressed as

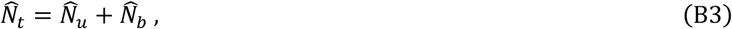

where 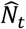 is the (normalized) total number of SF proteins in the cell, and 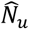 and 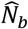 are the number of unbound and bound proteins, respectively. The number of bound proteins may be computed from:

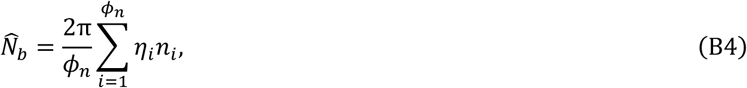

where *η*_*i*_ is the (normalized) cross-sectional SF concentration. We next consider the assembly of proteins into SFs. At steady state, the normalized SF concentration in direction *i* is given as:

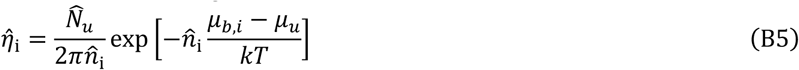

where *k* is the Boltzmann constant, and *T* is the absolute temperature. *μ*_*u*_ is the standard enthalpy of *n*^*R*^ unbound SF proteins, with *μ*_*u*_ = *μ*_*u*0_ + Δ*μ*_*u*0_*C*. The unbound proteins are affected by an activation signal *C* and form more readily into their bound states as the signal (e.g. concentration of unfolded ROCK) increases, and *μ*_*u*0_ is the standard enthalpy of the unbound SF proteins in the absence of a signal (*C* = 0) and Δ*μ*_*u*0_ the increase in the enthalpy of the unbound molecules at full signal activation (*C* = 1). At steady state we assume a continuous fully activated signal, i.e. *C* = 1. The second term on the right is the backward reaction rate for SF dissociation, with *μ*_*b,i*_ the standard enthalpy of *n*^*R*^ bound SF proteins, given as:

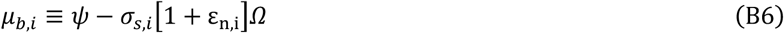

where *Ω* is the volume of *n*^*R*^ sarcomeres in a SF in an undeformed RVE, and *ψ* is the internal energy of *n*^*R*^ sarcomeres within a SF, given by:

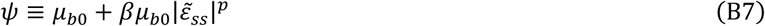

where *μ*_*b*0_ is the internal energy of *n*^*R*^ sarcomeres within a SF in their ground state, and *σ*_*s,i*_ is the tensile stress actively generated by a SF. In this paper we implement a steady state solution, hence the Hill tension-velocity relationship does not need to be considered as *σ*_*s,i*_ is necessarily equal to the maximum isometric tension *σ*_*max*_. Finally the 3D active stress tensor follows as:

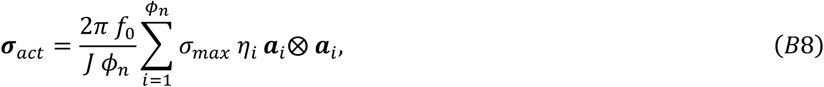

where *f*_0_ is the volume fraction of cytoskeletal proteins in the cell, and *J* is the determinant of the deformation gradient **F**.

#### Passive tissue model

In all regions of the tissue construct we consider the presence of an underlying isotropic material described by a simple neo-Hookean hyperelastic model, with the Cauchy stress

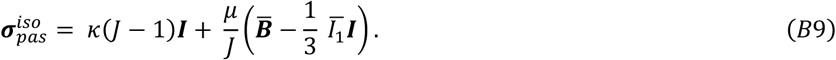

The first term on the right-hand side represents the hydrostatic stress contribution due to volumetric deformation, and the second term represents the deviatoric stress contribution due to isochoric deformation. ***B*** = ***FF***^*T*^ is the left Cauchy-Green tensor, with 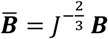, and ***I*** is the identity tensor. The first invariant ***I***_1_ is the trace of ***B***, with 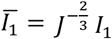, while *κ* and *μ* are the bulk modulus and shear modulus, respectively.

In order to account for dispersion of collagen fibres in the tissue construct, we implement a model adapted from the angular integration framework (Holzapfel and Ogden, 2015), whereby the fibre directions are discretely modelled. This allows for an exclusion of the mechanical contribution of all fibres under compression. We consider that fibres can exist in a large number of initially uniformly distributed discrete directions *ϕ*_*m*_ in a 3D sphere at each integration point of a finite element model (*ϕ*_*m*_ = 240 is found to provide a converged solution). This approach is adapted from our methodology for modelling cellular SF formation. The probability density function for dispersion is described by a von Mises distribution (Li et al., 2016), where the distribution factor in a given direction is expressed as

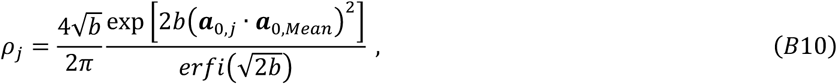

where *b* is a constant dispersion parameter, *erfi*(*x*) denotes the imaginary error function, ***a***_0,*Mean*_ is a unit vector indicating the mean fibre direction, and ***a***_0,*j*_ is a unit vector indicating one of *ϕ*_*m*_ directions. The distribution is normalized such that

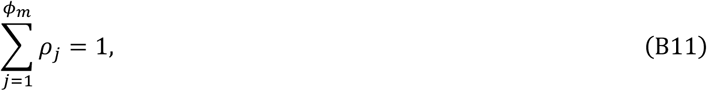

The collagen fibre stress is computed in all *ϕ*_*m*_ directions. The contribution of the dispersed fibres to the Cauchy stress tensor is then given as:

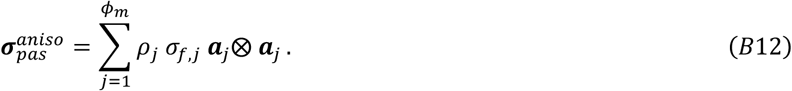

Finally, the total Cauchy stress for the tissue construct simulations is given by:

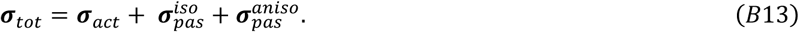

In the hydrogel simulations, the active cell contribution is excluded.

## Appendix C: Analysis of Active and Passive Transient Tissue and Hydrogel Behaviour

In this appendix we present a transient simulation in which we demonstrate the following: (i) Hydrogel stress relaxation is captured with a non-linear hyper-viscoelastic model; (ii) the transient evolution of active cell stress is captured using a novel cross-bridge cycling model that describes the key thermodynamics of actin-myosin interactions (which we have recently developed (McEvoy, Deshpande, McGarry, BMMB, 2019)). The transient model presented in this appendix is an extension of the steady state active model predictions that we have included in the main body of the paper.

A schematic of the model is shown in Figure C1. The model consists of the following components: (i) a non-linear hyper-viscoelastic hydrogel component, comprised of two nonlinear hyperelastic springs with a nonlinear viscous dashpot. (ii) a cell component consisting of an active contractile transient cross bridge cycling model (representing the active force generation of SFs), in parallel with a non-linear viscoelastic component (representing the passive cell contribution), comprised of a two nonlinear hyperelastic springs with a linear viscous dashpot.

**Figure C1.**
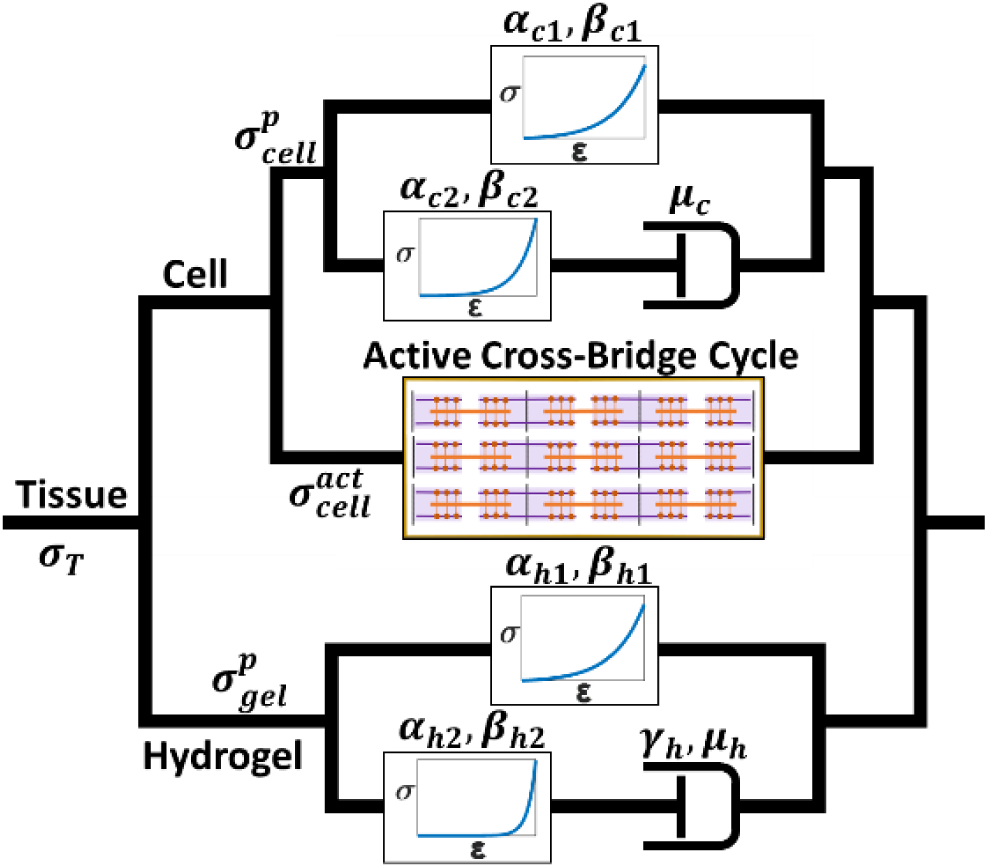
Schematic of the transient thermodynamic cross-bridge cycling cell model with non-linear hyper-viscoelastic hydrogel used to provide a preliminary interpretation of time dependent evolution in measured force during dynamic loading of uniaxial tissues.

### Model description

Total tissue stress, *σ*_*T*_, is given as

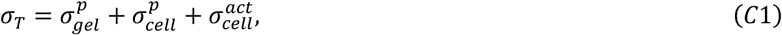

where 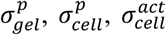 represent the passive hydrogel stress, the passive cell stress, and the active contractile cell stress, respectively. The constitutive law for the non-linear hyper-viscoelastic hydrogel component is given as:

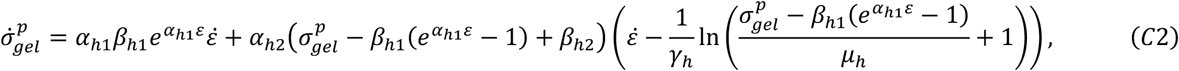

where *α*_*h*1_, *β*_*h*1_, *α*_*h*2_, *β*_*h*2_, *γ*_*h*_ and *μ*_*h*_ are model parameters. A similar non-linear viscoelastic constitutive law is used to represent the passive component of the cells:

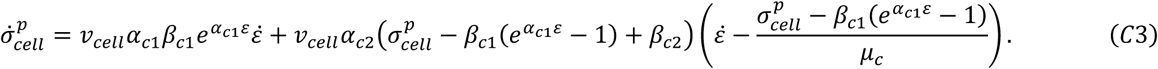

Again, *α*_*c*1_, *β*_*c*1_, *α*_*c*2_, *β*_*c*2_, and *μ*_*c*_ are model parameters. *v*_*cell*_ is the volume fraction of cells in the tissue.

We implement a thermodynamically based cross-bridge cycling law recently developed by McEvoy, Deshpande, McGarry (2019) to describe the dynamics of actin-myosin interactions within sarcomeres, resulting in transient active force generation. The rate of myosin-head attachment (cross-bridge formation) is given as

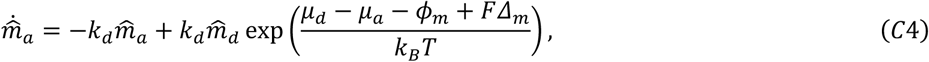

where 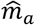 is the normalised number of attached myosin heads, *μ*_*d*_ is enthalpy of non-cycling myosin heads, *μ*_*a*_ is enthalpy of attached myosin heads, *ϕ*_*m*_ is stored elastic energy, *F*Δ_*m*_ is mechanical work performed by attached myosin heads, and *k*_*B*_*T* is Boltzmann’s constant and absolute temperature respectively. This process of cross-bridge formation results in active stretching of myosin tails, with the associated actively generated sarcomere tension *T*_*s*_ given as:

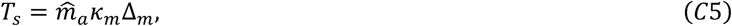

where *κ*_*m*_ is myosin tail stiffness, Δ_*m*_ is the myosin tail extension. This active generation of tension leads to the assembly of sarcomeres in parallel, such that:

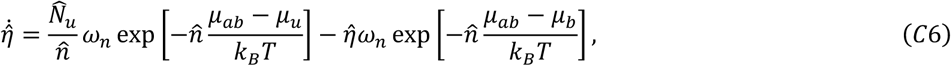

where 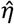 is a non-dimensional measure of parallel sarcomere formation, 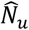 is the normalised total number of unbound sarcomeric proteins within a cell, 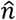 represents the normalised number of sarcomeres in series, *ω*_*n*_ is a collision frequency, *μ*_*u*_ is standard enthalpy of unbound proteins, *μ*_*b*_ is standard enthalpy of bound proteins, and *μ*_*ab*_ is an activation barrier, Vigliotti et al. (2015). This law leads to the computation of the active cell stress, given as

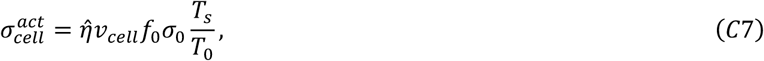

where the parameter *f*_0_ is the volume fraction of sarcomere proteins within the cell. The parameter *σ*_0_ is the isometric cell stress at zero shortening velocity. *T*_0_ is the computed sarcomere tension at zero shortening velocity.

### Model Boundary conditions and Parameters

The model is subjected to dynamic loading between 10% and 20% stretching at a frequency of 1 Hz. As shown in Figure 1D of the main paper, a sinusoidal wave form is applied, both in experiments and in simulations. Calibrated model parameters for the non-linear hyper-viscoelastic hydrogel model are as follows: *α*_*H*1_ = 4, *β*_*H*1_ = 0.025 *kPa, α*_*H*2_ = 15, *β*_*H*2_ = 0.012 *kPa, γ*_*H*_ = 1.1 *s, μ*_*H*_ = 120 *kPa*. Calibrated model parameters for the non-linear viscoelastic passive cell component are as follows: *α*_*C*1_ = 4, *β*_*C*1_ = .02 *kPa, α*_*C*2_ = 12, *β*_*C*2_ = .047 *kPa, μ*_*C*_ = 43000 *kPa s, v*_*cell*_ = 0.0049. All simulations are reported for cells at body temperature T = 310 K (∼ 37° C). Values implemented within cross-bridge cycle models are consistent with those given by McEvoy et al. (2019). The stroke distance is taken as *l*_*S*_ = 6.5 *nm*, the maximum allowable stretch during lengthening 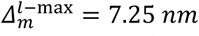, and the myosin tail stiffness *κ*_*m*_ = 1.75 *pN*/*nm*. With the sarcomere length is taken as *L*_*sarc*_ = 400 *nm*, and *t*_*s*_ = .001 *s* as the cross-bridge duty cycle times. The calibrated parameters *k*_*d*_ = 4 *Hz, γ* = 3, and *ε*_*s*_ = 0.225 with the time constant for signal decay as *τ* = 0.05 *s*. The enthalpies of attached and unattached cross-bridges, or bound and unbound SF proteins are taken as *μ*_*d*_ = 5 *k*_*B*_*T, μ*_*a*_ = 16 *k*_*B*_*T, μ*_*u*0_ = Δ*μ*_*u*0_ = 8*k*_*B*_*T, μ*_*b*0_ = 14*k*_*B*_*T*, and *μ*_*ab*_ = 20*k*_*B*_*T*. The remaining parameters for the SF framework are confined within ranges reported by Vigliotti et al. (2015) and McEvoy et al. (2019), with Ω = 10^−7.1^ *μm*^3^, *β* = 1.2, *p* = 2, and *f*_0_ = 0.011. Remodelling rates are calibrated to *ω*_*n*_ = 6 *Hz* and *α*_*n*_ = 1.2 *mHz*. The maximum isomeric stress *σ*_*iso*_ = 240 *kPa* is consistent with a wide range of measurements on muscle fibres (Lucas et al. 1987).

### Model Results and Comparison with Experimental Measurements

In Figure C2 we demonstrate strong agreement between model predictions and experimental measurements, both for the hydrogel material (which is modelled using just the lower non-linear hyper-viscoelastic component of the model (Figure C1)), and for the active tissue (which consists of both the cell component and the hydrogel component).

**Figure C2.**
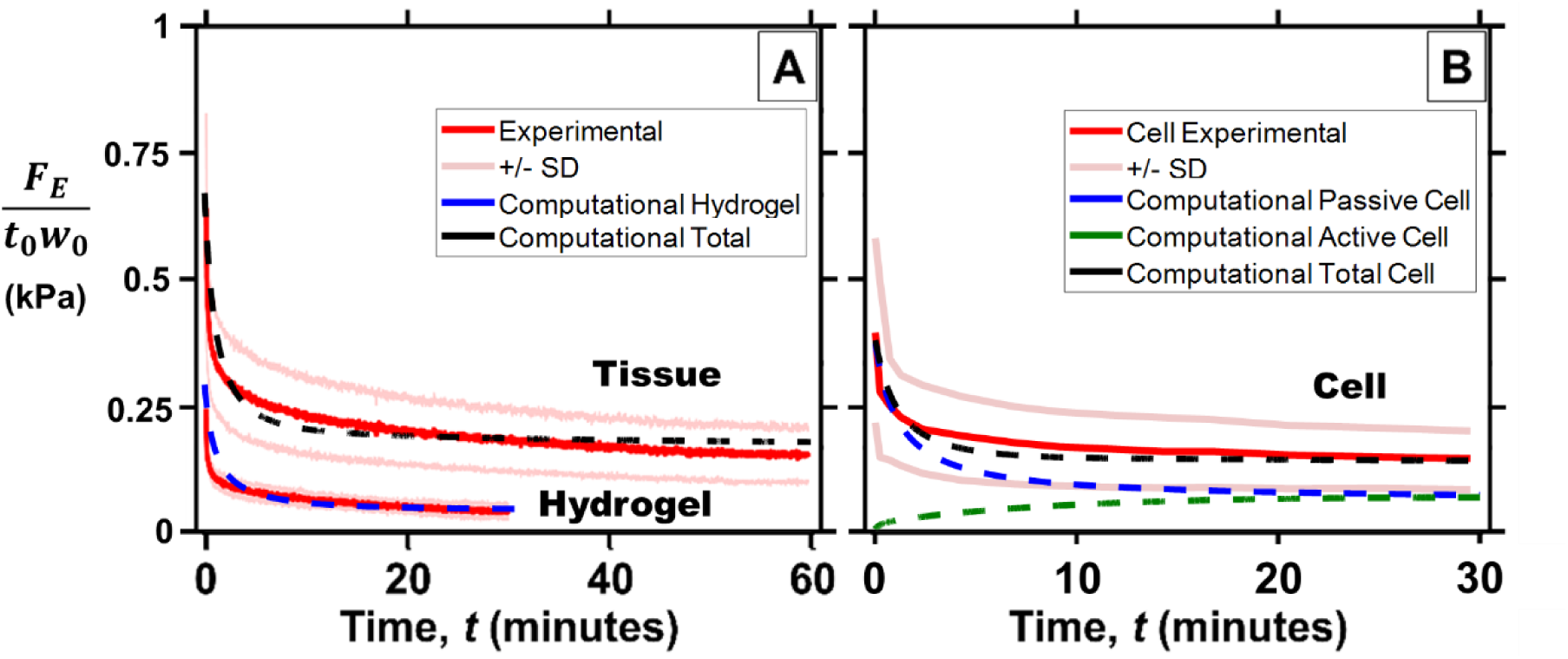
The maximum stress measured from cyclic deformation of: (A) Collagen tissues and hydrogels (Experimental mean (red curve) ± standard deviation (pink curves)) and (B) Cytoskeletal cell stress components (Interpolated experimental mean (red curve) ± standard deviation (pink curves)).

Key results are summarised as follows:

- Figure C2(A) demonstrates that the non-linear hyper-viscoelastic model accurately captures the stress relaxation of the hydrogel material.
- Figure C2(A) demonstrates that the full model (shown in Figure C1) (consisting of the transient sarcomere active cross-bridge cycling component, the passive non-linear viscoelastic cell component, and the non-linear hyper-viscoelastic hydrogel component) accurately captures the experimentally measured transient changes in tissue forces.

Figure C2(B) shows that the cell contribution of the model accurately captures the experimentally estimated cell contribution (nominal tissue stress minus nominal hydrogel stress). The model results suggest that the steady state cell contribution consists of ∼60% actively generated stress and ∼40% passive stress.

